# Despite the odds: formation of the SARS-CoV-2 methylation complex

**DOI:** 10.1101/2022.01.25.477673

**Authors:** Alex Matsuda, Jacek Plewka, Yuliya Chykunova, Alisha N. Jones, Magdalena Pachota, Michał Rawski, André Mourão, Abdulkarim Karim, Leanid Kresik, Kinga Lis, Igor Minia, Kinga Hartman, Ravi Sonani, Grzegorz Dubin, Michael Sattler, Piotr Suder, Paweł Mak, Grzegorz M. Popowicz, Krzysztof Pyrć, Anna Czarna

## Abstract

Coronaviruses protect their single-stranded RNA genome with a methylated cap during replication. The capping process is initiated by several nonstructural proteins (nsp) encoded in the viral genome. The methylation is performed by two methyltransferases, nsp14 and nsp16 where nsp10 acts as a co-factor to both. Aditionally, nsp14 carries an exonuclease domain, which operates in the proofreading system during RNA replication of the viral genome. Both nsp14 and nsp16 were reported to independently bind nsp10, but the available structural information suggests that the concomitant interaction between these three proteins should be impossible due to steric clashes. Here, we show that nsp14, nsp10, and nsp16 can form a heterotrimer complex. This interaction is expected to encourage formation of mature capped viral mRNA, modulating the nsp14’s exonuclease activity, and protecting the viral RNA. Our findings show that nsp14 is amenable to allosteric regulation and may serve as a novel target for therapeutic approaches.

## Introduction

The coronaviral genome is of positive polarity and serves as a substrate for the translational machinery of the cell after its release to the cytoplasm. The first and only product of the genomic mRNA translation is a large and non-functional 1a/1ab polyprotein. The polyprotein maturates by autoproteolytic processing carried out by two viral SARS-CoV-2 proteases – main protease, M^pro^ and papin-like protease, PL^pro^. This leads to the generation of a set of non-structural proteins (nsp), which are responsible for the viral replication process and remodeling of the intracellular environment. Once nsps reshape the cell to form a viral factory, the genomic RNA is copied and a set of subgenomic mRNAs is produced in a peculiar, discontinuous transcription process. These subgenomic mRNAs are monocistronic and serve as templates for the production of structural and accessory proteins required for the formation, assembly and release of progeny viruses^1^. The activity of particular nsps has been previously described, showing the complex network of interactions of multi-functional components. However, their coordinated action has not been fully understood.

Works by Gao *et al*.^2^, Yan *et al*.^3^, Wang *et al*.^4^, and Kabinger *et al*.^5^ shed light on the scaffold of the replication complex formed by nsp12 (polymerase) and two co-factors nsp7 and nsp8, that create functional machinery able to replicate the viral RNA. Next, an extended elongation complex was described, where nsp12/7/8 is accompanied by nsp13 helicase. This complex is suggested to serve as the basic replication module^6^.

Coronaviruses are known for their large genomes, requiring high-fidelity replication to maintain their integrity. While the SARS-CoV-2 nsp12 polymerase is highly processive, it is error-prone and does not provide sufficient fidelity. It has been previously demonstrated that a proofreading system is encoded in coronaviral genomes^7-9^. Nsp14 carries an N-terminal exonuclease (ExoN) domain that serves in this role. ExoN is a member of the DEDDh exonuclease superfamily and exhibits 3′-5′ exonuclease activity, removing incorrectly incorporated nucleotides from the 3′ terminus of the newly formed RNA. ExoN has additionally been proposed to play a role during discontinuous replication of coronaviruses. Nsp14 has been shown to associate with the replicatory complex, with nsp10 as a co-factor, modulating and enhancing nsp14 exoribonuclease activity^7-9^.

Apart from the supporting role in replication, nsp14 has a second important function. It takes part in cap formation after genome copying is finalized^7^. Capping of viral mRNAs is essential for their function and integrity. It enables translation initiation and protects viral mRNA from recognition as foreign by cellular sensors, thereby preventing the induction of innate immune responses^10^. Cap formation is a tightly regulated process consisting of four consecutive enzymatic reactions. First, nsp13 triphosphatase removes the γ-phosphate of the 5′-triphosphate end (pppA)^11,12^; next, nsp12 guanylyltransferase (the nidovirus RdRp-associated nucleotidyltransferase, NiRAN domain)^3^ transfers GMP to the 5′ phosphate to form the core structure of the cap (GpppA); the GpppA is methylated at the N7 position by the nsp14 N7-methyltransferase domain (^7Me^GpppA); subsequently, ribose in the first ribonucleotide is methylated at the 2′-O-position by the nsp16 2′-O-methyltransferase^13-15^. Capping is regulated by nsp9, which binds nsp12 near the NiRAN active site^16^. The described process results in a functional cap (^7Me^GpppA_2′OMe_), completing the genome replication. While there is a good structural understanding of the interactions between nsp12, nsp13, and nsp14 during replication, nsp16 remains an orphan, and its methylation process is not fully understood.

Here, we studied the interaction between the two methyltransferases encoded in the genome of the coronavirus–nsp16 and nsp14. Both protteins bind to nsp10 and both are required for complete methylation. Therefore, their spatial proximity mediated by nsp10 would appear as beneficial for the methylation efficacy. Prior *in silico* analysis of structural information on nsp10/14 and nsp10/16 complexes suggested that simultaneous binding of both exonucleases to nsp10 should be impossible due to steric hindrance^17^.Interestingly, both nsp14 and nsp16 interact with nsp10 by typical, well-defined protein-protein interfaces containing deep-reaching lipophilic residues and solvent-protected hydrogen bonds. The element that causes steric overlap between nsp14 and nsp16 is a peculiar N-terminal “lid” domain of nsp14. This lid is mostly devoid of secondary structure and lacks characteristic protein-protein interaction complementarity. Interestingly, recent structures of nsp14 without nsp10 show massive rearrangement of the lid region confirming its structural flexibility^18,19^. This prompted us to postulate that a local structural rearrangement of the nsp14 N-terminal lid region is therefore required and possible, thus facilitating the heterotrimer complex formation.

## Results

### Biochemical evidence of heterotrimer formation

Nsp10, nsp14, and nsp16 were co-expressed in *E. coli* and purified. A single peak was obtained in size exclusion chromatography (SEC), indicating that all three proteins co-migrate (Fig 1a). Because no prior crosslinking was used, this result suggests the formation of a heterotrimer complex. Sodium dodecyl sulphate-polyacrylamide gel electrophoresis (SDS-PAGE) analysis of the peak containing the putative complex, together with mass spectrometry identification of components, indicates the presence of all three proteins (Fig. 1b). We further show that the putative complex migrates as a single major band in native PAGE (Extended Data Fig. 1a). When the band was excised from the gel and analyzed by SDS-PAGE, three bands were identified, corresponding in molecular weight to nsp14, nsp16, and nsp10 (Extended Data Fig. 1b). MS analysis of proteins contained in the major band derived from the native PAGE resulted in the identification of all three components of interest (Extended Data Table 1), further suggesting heterotrimer complex formation. LC-MS additionally allowed the assessment of the stoichiometry of the complex. By analyzing the signals at 254 and 280 nm, with further MS-based identification of the proteins under each UV chromatographic peak, stoichiometry was consistently established at 1.2:1:1 (nsp10:nsp14:nsp16) (Extended Data Fig. 2 and Extended Data Table 2).

**Fig. 1.**
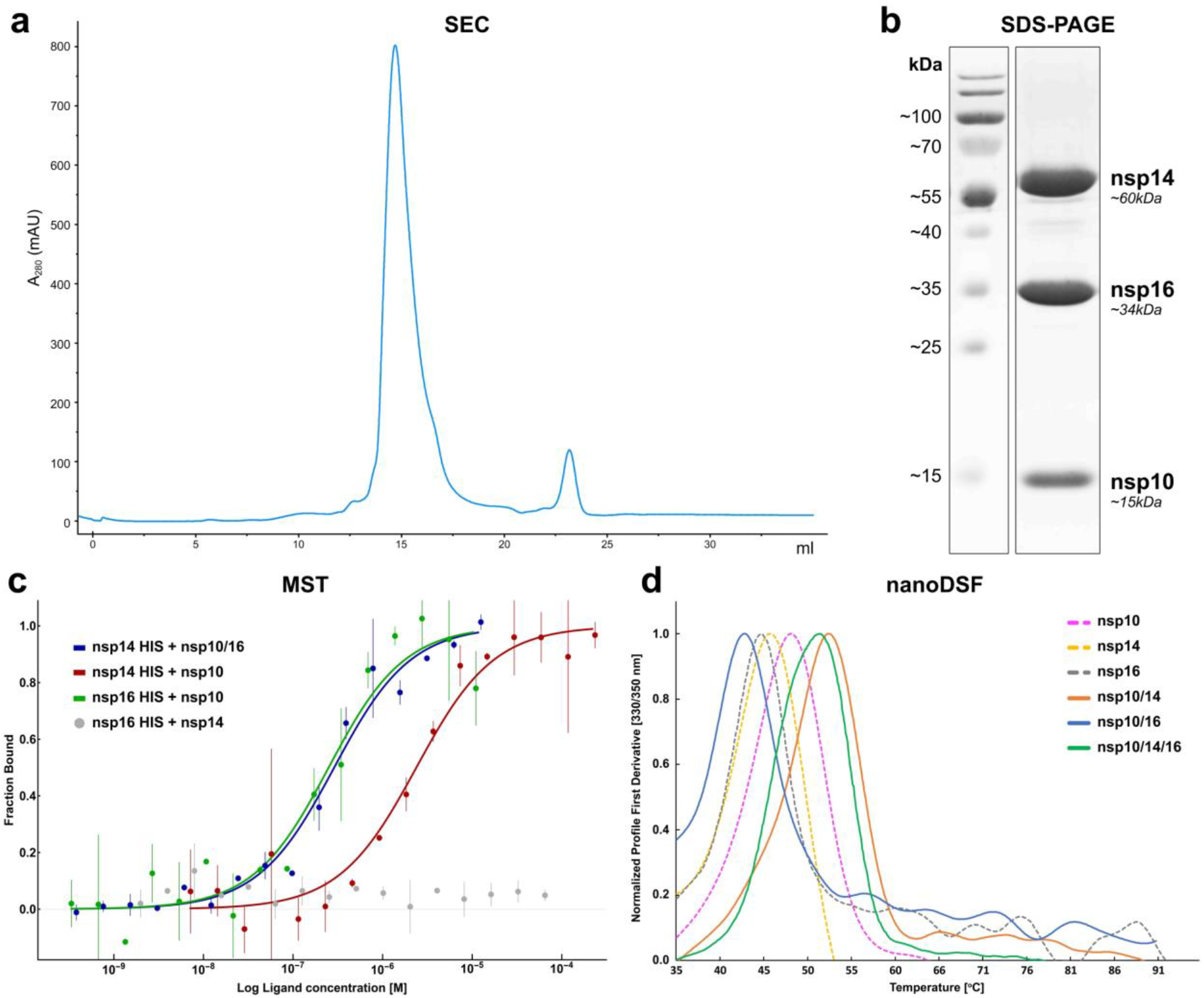
In vitro formation of the nsp10/14/16 heterotrimer. **a**, SEC chromatogram showing co-migration of nsp10, nsp14, and nsp16 proteins in a single peak at ∼15 mL. **b**, SDS-PAGE analysis of the major peak from panel a. Three major proteins corresponding in size to nsp10, nsp14, and nsp16 (Mw of 15, 60, and 33.5 kDa, respectively) are present in roughly equal quantities. **c**, MST analysis of the intra-complex affinities. Experimental data are represented as mean (dots) with error bars. Binding model fit is represented as a solid line. **d**, The melting profiles of heterotrimer complex and complex components, determined using nanoDSF.

The kinetics of the heterotrimer complex formation were assessed by MicroScale Thermophoresis (MST). Nsp14 and nsp16 were separately expressed as histidine-tagged constructs and labeled with a high-affinity His-tag specific fluorophore dye. When labeled nsp14 was titrated with unlabeled nsp10, a dose-dependent increase in thermophoretic signal was observed, indicating an interaction with a K_d_ of 2.4 ± 0.2 µM, which is in the agreement with the previously reported affinity with a K_d_ of 1.1 ± 0.9 μM^20^ (Fig. 1c). A comparable effect^21^ was observed when labeled nsp16 was titrated with unlabeled nsp10 with a K_d_ of 0.24 ± 0.01 µM. Labeled nsp14 did not directly interact with unlabeled nsp16, but when labeled nsp14 was titrated with unlabeled nsp10/16 complex, a dose-dependent increase in thermophoretic signal was observed (Fig. 1c), which was interpreted as the heterotrimer complex formation. Fitting the experimental data allowed the determination of the K_d_ characterizing the interaction at 0.28 ± 0.01 µM.

To further characterize the heterotrimer complex and its components, we analyzed thermal denaturation profiles using nanoDSF. Each of the individual system components (nsp10, nsp14, and nsp16) was characterized by a single characteristic denaturation temperature (Fig. 1d), indicating that structures of the functional domains (if any) within particular components collapse in a coordinated manner upon temperature increase. Nsp16 was least temperature stable (T_m_=45°C), while nsp10 (T_m_=50°C) was most stable. However, the differences in melting temperatures were small, with all components demonstrating overall similar stability. Nsp14 was stabilized by the complex formation with nsp10 (nsp10/14 complex: T_m_=55°C), while nsp10 binding did not significantly affect the thermal stability of nsp16 (nsp10/16 complex: T_m_=45.5°C). The heterotrimer is characterized by a single sharp thermal denaturation peak with a characteristic melting temperature of 51.7°C. This temperature, distinctly different from that of any of the individual or binary-complexed components, gives a further evidence for the heterotrimer complex formation.

### Functional consequences of nsp10/14/16 heterotrimer complex formation

GpppA is methylated at position N7 by nsp14 N7-methyltransferase domain, yielding N^7^MeGpppA^22^; SAM is used as a donor of the methyl group (Fig. 2a). Here, we tested whether the heterotrimer complex formation affects nsp14 catalyzed methylation. To follow the N7-methyltransferase activity of nsp14, we used an indirect assay to monitor changes in the level of a second reaction product, SAH, by HTRF. In the absence of one of the substrates (SAM or RNA) or the tested enzyme (negative controls), there was no activity validating the experimental setup (Extended Data Fig. 3a). Nsp14 showed no preference for the nascent nucleotide methylating both GpppG and GpppA, as demonstrated by the production of equal levels of SAH when using either nucleotide as a substrate (Fig. 2b). The binary complex of nsp10/14 and the ternary complex N7-methyltransferase activities were comparable to that of nsp14 alone, indicating that binding to nsp10 or the heterotrimer complex formation had no significant influence on the N7-methyltransferase activity of nsp14. N7-methyltrasnferase activity was also not reported for N7-methylated substrate (O4 oligo - 5’ (N7-MeGppp) ACA UUU GCU UCU GAC 3’). Nsp16, on the other hand, presenting 2′-O-methyltransferase activity in the complex with its obligatory partner nsp10 does not show a signal on non-methylated substrates and modetare activity on the O4-oligo substrate. Similarly to N7-methyltransferase activity, also 2′-O-methyltransferase activity is not affected by the heterotrimer complex formation. As expected, the pan-methyltransferase inhibitor sinefungin^23^ halted methyltransferase activities of all repoted here proteins. We have also tested if the nsp14 exoribonuclease (ExoN) activity affects methyltransferase activity by mutating the ExoN binding site. However, no difference in N7-methyltrasnferase activity was observed, decoupling those two activities from each other (Extended Data Fig. 3b). Alongside N7-methyltransferase activity, nsp14 harbors an ExoN domain characterized by exoribonuclease activity; an activity essential for proofreading during virus replication. However, *in vitro*, nsp14 is characterized by the high processivity and non-specifically degrades nucleic acids^17,24,25^. As the nuclease activity must be tightly controlled, we assessed whether the complex formation regulates this process. We evaluated the binding and nuclease activity of nsp10/14 and nsp10/16 compared to nsp10/14/16 with two synthetic RNAs of CoV-RNA1-G 5’-GGGGGGGGGGGCGCGUAGUUUUCUACGCG-3’ and CoV-RNA1-A 5’-AAAAAAAAAAACGCGUAGUUUUCUACGCG-3’.

**Fig. 2.**
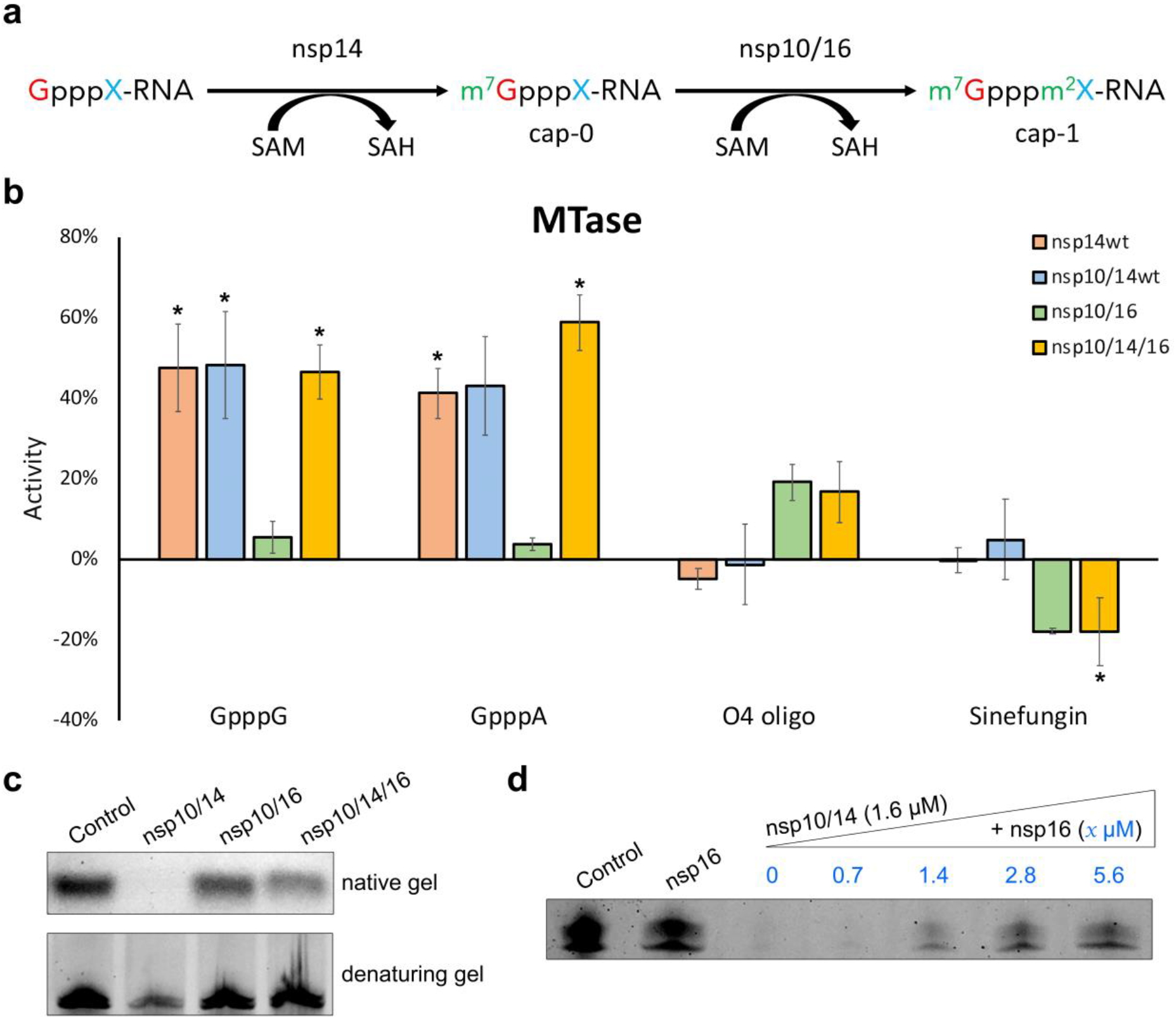
The formation of nsp10/14/16 heterotrimer modulates the ribonuclease, but not methyltransferase processivity. **a**, Schematic view of mRNA methylation. X represents a nascent nucleotide that could be adenine (A) or guanine (G). m^7^ represents methylation of the first guanine at position N7 by nsp14. m^2^ indicates 2′-O methylation of the nascent mRNA nucleotide. **b**, Modulation of the methyltransferase activity by the protein partners. The results were normalized using the SAH calibration curve and denoted µM of methylated product. All experiments were performed in duplicate. Average values with error bars (SD) are shown; * p < 0.05 in comparison to the condition with no activity. **c**, Analysis of RNA binding (native gel) and degradation (denaturing gel) potential of indicated nsp complexes. **d**, Determination of stoichiometry of the heterotrimer complex limiting the exonuclease activity of nsp14. Titration of nsp10/14 (1.6µM) with indicated amounts of nsp16 (blue). (c, d) CoV-RNA1-A was used as a substrate. Similar results are obtained for CoV-RNA1-G (Extended Data Fig. 4).

Addition of nsp10/14 and nsp10/16 (equimolar at 1.6µM) results in the retardation of the RNA relative to an RNA only control, thus indicating a binding interaction between the RNA and protein heterodimers (Fig. 2c native gel, Extended Data Fig. 4). Interestingly, upon addition of nsp10/14/16 (equimolar at 1.6 µM), we observed increased retardation of the RNA-heterotrimer protein complex relative to the nsp10/14 and nsp10/16 heterodimer complexes (Fig. 2c, native gel). The observed broadened band suggests that nsp10/14/16 forms a tetrameric complex with each of the RNA substrates.

As expected, addition of nsp10/14 results in the degradation of the RNA substrates (as indicated by the decreased intensity of the RNA band in the presence of protein relative to the RNA only control, observed both by native and nondenaturing gel analysis) (Fig. 2c, denaturing gel). Interestingly, we observed that the nuclease activity of nsp10/14 is reduced for the nsp10/14/16 heterotrimer, suggesting that nsp14 nuclease activity is modulated by nsp16 (Fig. 2c, denaturing gel). To assess this, we added increasing amounts of nsp16 to preformed nsp10/14 CoV-RNA1-A RNA complex and monitored the degradation of the RNA. Upon increasing the concentration of nsp16, the degradation activity of nsp14 is reduced, starting at a near 1:1 ratio (Fig. 2d). This effect is unlikely to be caused by the protective effect of RNA sequestering by the nsp16 as the RNA-nsp10/16 affinity is reported to be in high micromolar regime^26^ (∼100 µM) and yet protective functions of nsp16 appear at much lower protein concentrations equal to the nsp10/14 complex concentration. Recent crystallographic data suggest that nsp14 exonuclease domain is controlled by lid rearrangement caused by nsp10 binding^18^. In combination with our binding shift and degradation assays, this data suggests that binding of nsp10/16 to nsp14 causes additional allosteric change that inhibits unwanted exonuclease activity in favor of the methyltransferase one.

### Lid hypothesis explains the nsp10/14/16 heterotrimer complex formation

A number of high-resolution crystal structures are available for the nsp10/14 and nsp10/16 complexes. Overlay of the structures by a common component (nsp10) demonstrates that fifty N-terminal amino acids of nsp14 overlap with nsp16 at the surface of nsp10. The significant steric clash produced by this overlap precludes the heterotrimer complex assembly, which is mediated by the concomitant interaction of nsp14 and nsp16 with nsp10 (Fig. 3a-c). As such, the formation of the heterotrimer complex would require a significant structural rearrangement within either nsp14 or nsp16. Analysis of the interactions between nsp10, nsp14 and nsp16 suggests that the former binding surface contains a weakly interacting component – the N-terminal region of nsp14 (for rigorous analysis, see Discussion). We hypothesize that within the nsp10/14/16 heterotrimer complex, nsp16 displaces the N-terminal “lid” of nsp14 at the interface with nsp10.

**Fig. 3.**
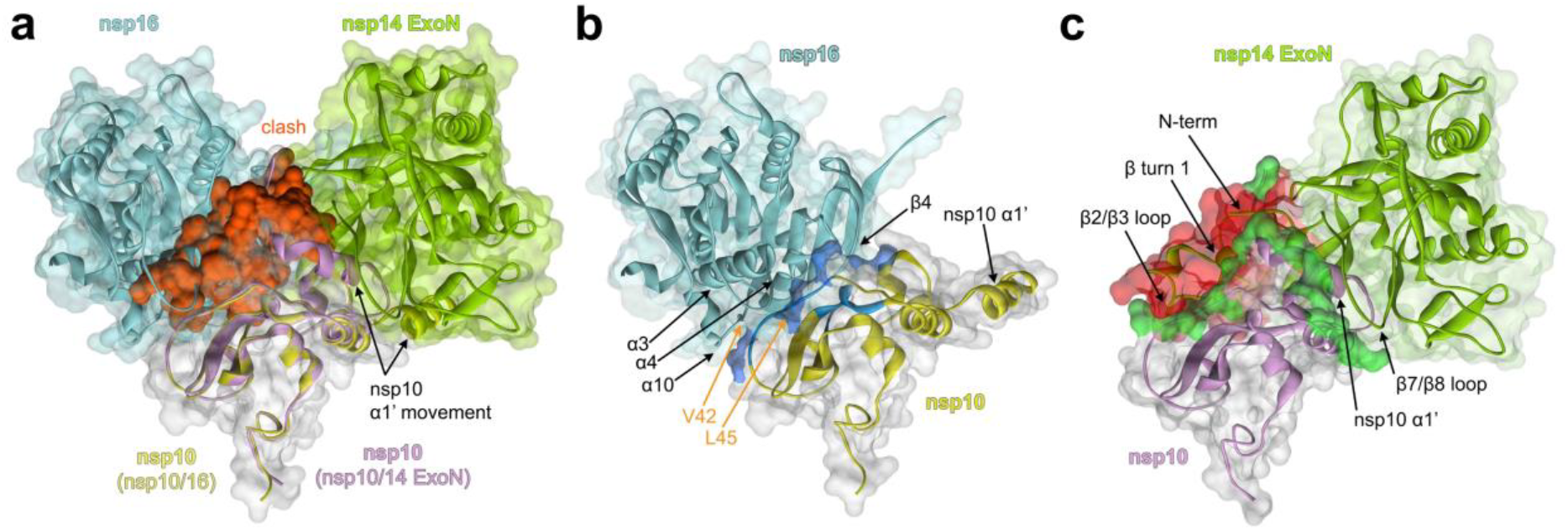
In silico modelling of the nsp10/14/16 complex. **a**, Nsp10-centered alignment of nsp10/16 (PDB ID: 6WVN) and nsp10/14 ExoN domain (PDB ID: 7DIY). Nsp16 is shown in cyan, and the associated nsp10 is shown in yellow. nsp14 ExoN domain is shown in green and the associated nsp10 in purple. The structural clash between the nsp14 ExoN and nsp10/16 surfaces is shown in red. **b**, The interface between nsp10 and nsp16. The strong hydrophobic interaction between nsp10 and nsp16 is shown in blue. The nsp10 residues V42 and L45 are shown as sticks. **c**, The interface between nsp10 and the nsp14 ExoN. The hydrogen bonds between nsp10 and nsp14 are shown in green. All panels: Different orientation of nsp10 α1’-helices in respective complexes with nsp16 and nsp14 is shown. Arrows indicate major structural features constituting the interface.

To evaluate the above hypothesis, we created a “lid”-truncated mutant of nsp14 (nsp14Δ) missing the initial 50 amino acids and evaluated its interaction with nsp10. Nsp14Δ still formed a complex with nsp10, characterized by a K_d_ of 1.5 µM. This value did not differ significantly from that characterizing the complex involving full length nsp14 (K_d_=2.4 µM). The above data indicate that the interactions of the N-terminal region of nsp14 with nsp10 do not contribute significantly to the affinity of either component, indirectly supporting the “lid” rearrangement hypothesis in the heterotrimer complex formation.

### Structural characterization of the heterotrimer complex

Nsp14, nsp10/14, nsp10/16 and nsp10/14/16 were characterized by SEC-SAXS^15^. The retention times of tested biomolecules correlate with theoretical molecular weights, assuming 1:1(:1) molar ratios^16^ (Extended Data Fig. 5). Guinier analysis of scattering profiles established that the largest radii of gyration (R_g_) characterized the heterotrimer complex (40.3±1.0 Å). Nsp14 and the nsp10/14 complexes were characterized by significantly smaller R_g_s (28.0±0.5 and 30.1±1.8 Å, respectively); the nsp10/16 complex was characterized by the smallest (R_g_ = 21.0±1.5 Å) (Extended Data Table 3). This data corresponds well with the expected molecular weights of tested complexes at 1:1:1 stoichiometry, further supporting the stoichiometry of the heterotrimer complex.

The estimated molecular weights of a scatterer differ from the expected values, perhaps as the result of the flexibility of the tested system (SAXS signal averages all conformations). Nonetheless, the relative values follow the expected pattern, with nsp10/16 characterized by the lowest and the nsp10/14/16 heterotrimer complex by the highest molecular weight, as determined by SAXS.

When reconstructed in real space using the indirect Fourier transform using software GNOM, the scattering profiles present roughly Gaussian shapes with significant tailing for nsp10/14/16 and nsp10/14, indicating an elongated globular nature for the protein complexes, with peaks overlapping with radii of gyration obtained using Guinier analysis. Calculated maximal distances within scatterers support the trend established above, with nsp10/16 constituting the smallest complex at 80.0 Å, nsp14 at 95.8 Å, nsp10/14 at 122.0 Å, and the nsp10/14/16 at 140.0 Å (longest axis).

Molecular envelopes which best represent the scattering profiles were calculated using DAMMIF software^27^ (Fig. 4). Crystal structures of nsp10/16, nsp14 and nsp10/14 and a model of the nsp10/14/16 heterotrimer complex, created assuming the “lid” hypothesis, were fitted into the envelopes using the SUPCOMB software. The crystal structures of nsp14 and nsp10/14 fit the molecular envelopes poorly, suggesting that these two remain flexible in solution (Fig. 4a and Fig. 4b). The structure of nsp10/16 fills the envelope tightly, suggesting complex rigidity in solution. The initial model of the heterotrimer complex already filled the envelope relatively well and was further optimized via normal mode analysis using the SREFLEX software^28^. SREFLEX rotates and translates rigid body domains of the input model within the constraints of flexible loops, optimizing the fit to the experimental scattering curve. The resulting heterotrimer model was characterized by a value of 1.08 for goodness-of-fit to the experimental SAXS data, suggesting a very likely solution. The model fits the envelope tightly, suggesting the rigidification of nsp14 structure upon the heterotrimer complex formation (comparing to nsp10/14 envelope fit). The decomposition of the nsp10/14/16 scattering profile into volume fractions calculated from the binary complexes nsp10/14, nsp10/16 crystal structures or extracted individual protein in the software Oligomer suggest that the signal can be devided 1:1 into nsp10/16 and nsp14 (Extended Data Table 3). This furter implies that nsp10/16 interface within the nsp10/14/16 is retained, while it is nsp14 that undergoes structural rearrangements upon the heterotrimer complex formation.

**Fig. 4.**
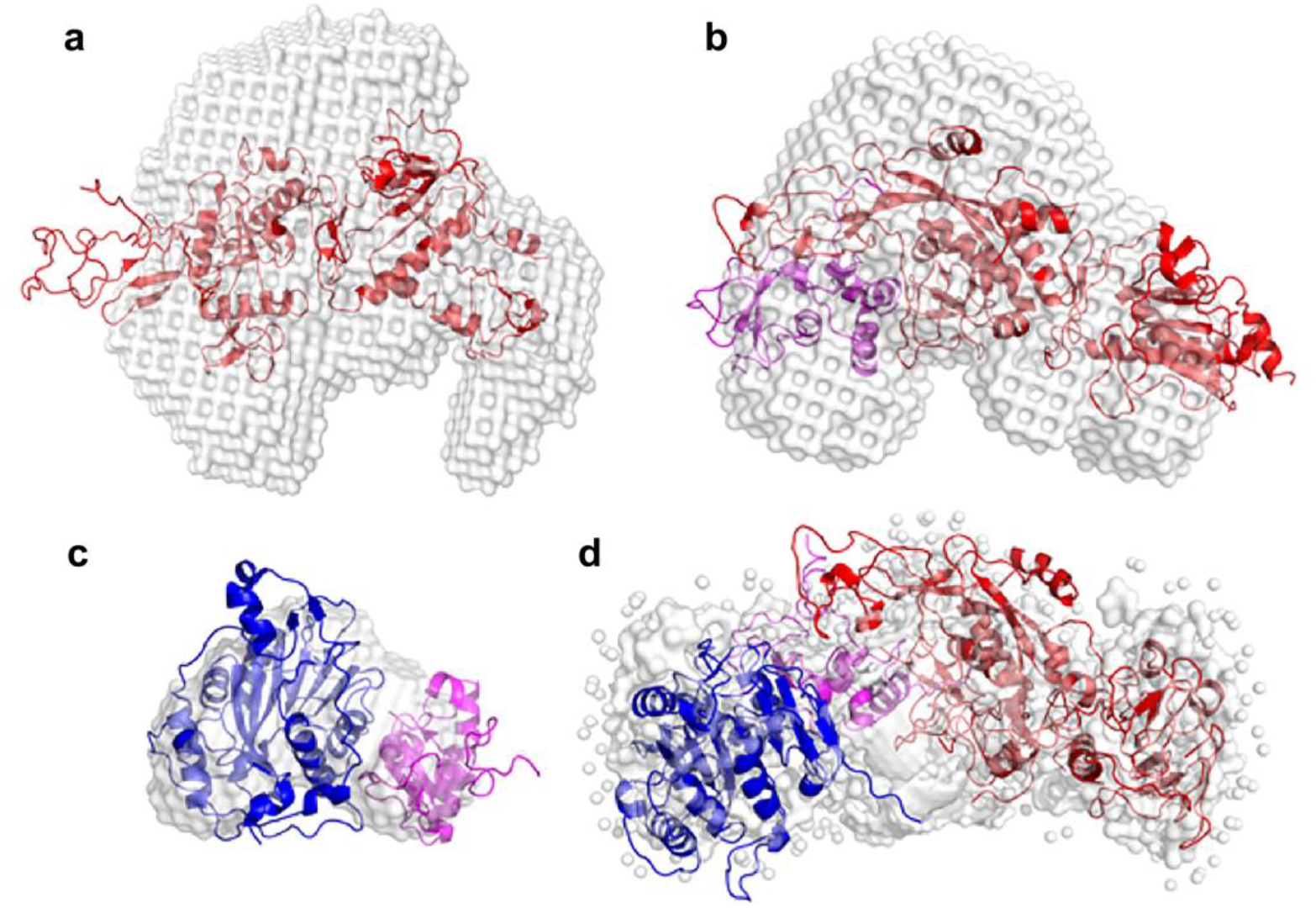
Molecular envelopes representing the experimental SAXS scattering profile of nsp14 (**a**), nsp10/14 (**b**), nsp10/16 (**c**), nsp10/14/16 (**d**). Overlaid are best fits of crystallographic / theoretical models of relevant complexes. Color coding: nsp10 in magenta, nsp14 in red, nsp16 in blue, molecular envelopes in grey.

The heterotrimer complex formed by nsp10, nsp14 and nsp16 was further characterized by transmission electron microscopy. Negatively stained samples of a heterotrimer complex formed from full-length components yielded a non-homogenous particle distribution, which precluded structural analysis (Extended Data Fig. 6). However, when nsp14 methyltransferase domain was truncated out of the structure leaving nsp14 ExoN, the particle distribution became more homogenous, allowing the convergent classification (Fig. 5a) and structural analysis. The reconstruction obtained from the negative-stained transmission electron micrographs at 20 Å resolution shows elongated particles with approximate dimensions of ∼ 10 × 5 × 4.5 nm (Fig. 5b). Necking is evident in the center of the particles, indicating that two larger structural components are connected by a component of a lower molecular weight (nsp10). The SAXS-derived structural model of the heterotrimer complex fits the experimental NS-TEM map reasonably well with CCmask/CCbox of 0.59/0.80. Nsp14 ExoN and nsp16 fit the two globular regions of the map, connected by a region with a density corresponding to nsp10. The central cavity suggested by SAXS data is visible in the electron microscope-derived map, indicating an overall match between the 3D reconstitutions obtained using each method.

**Fig. 5.**
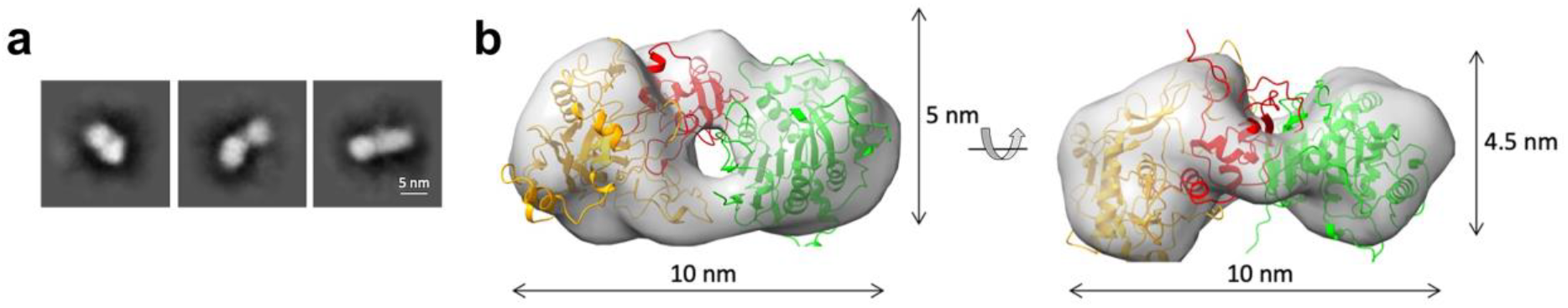
Transmission Electon Microscopy characterization of nsp10/14Δ/16 complex. **a**, Representative 2D-classes obtained by template-free 2D classification of particles picked from NS-TEM micrographs of nsp10/14/16 complex. **b**, Rigid body fit of SAXS-derived structure of the heterotrimer complex into NS-TEM-derived 3D reconstitution map. Nsp14, nsp10, and nsp16 are represented as orange, red, and green ribbon models, respectively. The 3D reconstitution map is shown as a transparent grey surface.

## Discussion

It was demonstrated earlier that nsp10 serves as a protein co-factor of both nsp14 and nsp16. One might logically expect that nsp10 could bring nsp14 and nsp16 together into a heterotrimer complex and that the kinetics and processivity of the process would be positively influenced by the spatial proximity of enzymes catalysing consecutive reactions in the capping pathway. The crystal structures available to date, however, suggest otherwise. The binding interfaces of nsp14 and nsp16 overlap at the surface of nsp10, suggesting that the heterotrimer complex is not feasible without a major structural rearrangement of either nsp14 or nsp16.

Careful analysis of existing data demonstrates that the nsp10/16 interface relies on a solid network of hydrophobic interactions mediated by a rigid central antiparallel β1-sheet of nsp10; the helices α2, α3 and α4; a coiled-coil region connecting helix α1 and the sheet β1 and α helices 3,4 and 10; as well as β-sheet 4 (Fig. 3b). In particular, Val42 and Leu45 of nsp10 are embedded into hydrophobic pockets formed by helices α3, α4, and α10 of nsp16.

Despite the fact that the interface between nsp10 and nsp14 buries a larger surface area compared to the nsp10/16 interface, the affinity characterizing the components of the former complex is almost an order of magnitude weaker than that of the latter. This prompted us to speculate that a significant part of the nsp10/14 interaction may not significantly contribute to affinity. The nsp10/14 interaction surface contains two major regions. Interactions within the N-terminal region of nsp14 involve primarily loops and other poorly structured regions, such as the N-terminal coil-coiled region that interacts with the α1’ helix of nsp10, β-turn 1, and loops between β-sheet 2/3 and β-sheet 7/8 (Fig. 3c). Many of those interactions are present within the region that would overlap with nsp16 if the complexes were aligned. Indeed, nsp14Δ retains nsp10 binding properties and is characterized by an affinity comparable to wild type nsp14, supporting the claim that the N-terminal region does not significantly contribute to interaction. This allowed us to hypothesize that the heterotrimer complex is formed when nsp16 displaces the N-terminal region (“lid”) of nsp14 at the surface of nsp10 (Fig. 6). Indeed, both nsp14 and nsp14Δ readily form a heterotrimer complex with nsp16 and nsp10.

**Fig. 6.**
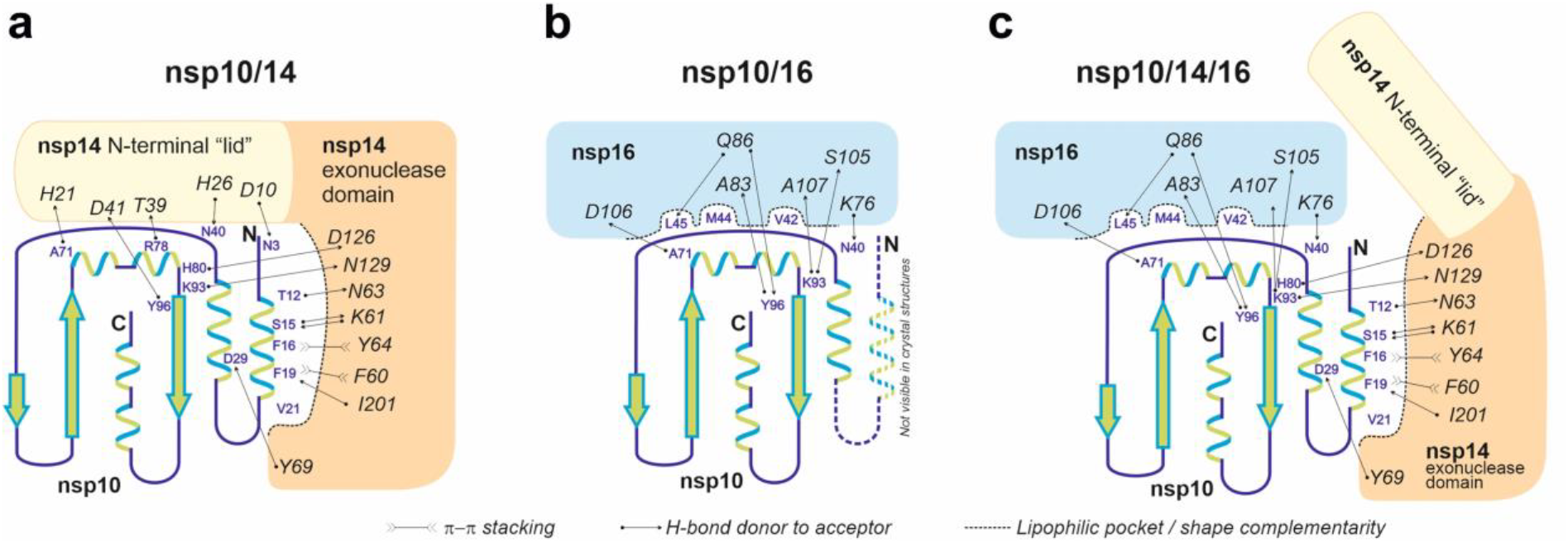
Schematic diagram of interactions guiding the affinities of nsp10, nsp14 and nsp16. **a**, The α1 helix of nsp10 provides several deeply buried lipophilic and π-stacking interactions with the exonuclease domain of nsp14 and the interaction may be described in terms of shape complementarity. In turn, the interaction of the N-terminal “lid” region (amino acids 1-50) of nsp14 with nsp10 shows poor shape complementarity and is characterized only by a low number of solvent-exposed hydrogen bonds. **b**, Nsp16 binds nsp10 at the site overlapping that involved in binding of the “lid” of nsp14, but nsp10/16 interaction is characterized by a well-developed interface involving deep lipophilic pockets and solvent-shielded hydrogen bonds. The α1 helix of nsp10 is not defined in the crystal structures of the nsp10/16 complexes (6W4H, 6YZ1), indicating it is flexible and not involved in binding. **c**, The formation of the nsp10/14/16 heterotrimer complex is accompanied by “lid” displacement and stabilization of the α1 helix.

The “lid” hypothesis is supported by low-resolution structural data provided in this work. The gyration radii and molecular weight of the heterotrimer complex derived from SEC-SAXS experiments are higher than the gyration radii and molecular weights of any of the components or binary complexes (Extended Data Fig. 5a). The molecular envelope derived *ab inito* from the scattering profile of the heterotrimer complex perfectly fits the model suggested by the “lid” hypothesis. Additionally, SAXS demonstrates that the heterotrimer complex formation stabilizes conformational flexibility of the nsp10/14 complex.

NS-TEM reconstruction of 3D volume characterizing the nsp10/14Δ/16 heterotrimer complex accommodates the SAXS-derived model with high confidence, further supporting the lid hypothesis. We were unsuccessful in our effort to use cryoEM to determine a high-resolution structure of the heterotrimer complex composed of full-length components due to low particle homogeneity. Low resolution reconstruction was nonetheless possible and again accommodated the SAXS derived model well (Extended Data Fig. 7), further supporting the structural arrangement of the components within the heterotrimer complex.

Overall, in this study we provide evidence that nsp14, nsp10 and nsp16 form a heterotrimer complex characterized by 1:1:1 stoichiometry, built around nsp10. The architecture of the complex follows the general arrangement previously observed in nsp10/16 and nsp10/14 complexes, but nsp16 displaces the “lid” (N-terminal) of nsp14 at the nsp10 surface (Fig. 6). The heterotrimer complex brings together two consecutive activities required for RNA cap formation (nsp14 associated N7-methyltransferase and nsp16 2′-O-methyltransferase), likely contributing to the processivity of the capping process. The heterotrimer complex formation does not, however, influence the methyltransferase activities. Further, the heterotrimer complex formation mitigates the 3′→5′ exonuclease activity of nsp14, preventing the excessive degradation of viral nucleic acid and allowing the complex to switch from the proofreading mode to the methylation mode.

## Materials and Methods

### Protein expression and complex purification

Constructs of nsp10, comprising amino acids 4254 – 4392, nsp14, comprising amino acids 5926 – 6452, and nsp16, comprising amino acids 6799 – 7096, of SARS-CoV-2 polyprotein 1ab optimized for expression in *E. coli* were ordered from GeneArt and subcloned into expression vector pETDuet-1. For the nsp14 catalytic mutant (np14-cat), D90 and E92 were replaced with alanines^29^. Plasmids were then co-transformed into *E. coli* strain BL21.

Transformed BL21 (DE3) *E. coli* cells were grown in Terrific Broth medium (TB; Bioshop), supplemented with 100 µg/ml of ampicillin (Sigma), at 37 °C overnight and used as a starter culture for the large-scale expression in TB. After the culture reached OD_600_=1.2 – 1.4, it was induced with 0.5 mM isopropyl-D-1-thiogalactopyranoside (IPTG; Sigma), and protein expression was carried out at 18 °C for 16 hours. Bacterial pellets were collected by centrifugation at 6’000 rpm for 10 min at 4 °C, resuspended in lysis buffer (50 mM Tris-HCl pH 8.5, 300 mM NaCl, 5 mM MgCl_2_, 5% v/v glycerol, 5 mM β-mercaptoethanol) and disrupted by sonication at 80% amplitude for 15 min at 10°C (3s, 3s pulse). The lysed sample was clarified by centrifuging for 1 h at 25’000 rpm at 4°C. The supernatant was collected and incubated with 2 ml of Ni-NTA Agarose (Jena Bioscience) pre-equilibrated with the lysis buffer for 2 hours at 4°C. Purification was carried out in a gravity-flow column. Resin was washed with 50 BV (bed volume) of buffer A (50 mM Tris-HCl pH 8.5, 300 mM NaCl, 5 mM MgCl_2_, 5 mM β-mercaptoethanol, 10 mM imidazole) and 20 BV of buffer B (50 mM Tris-HCl pH 8.5, 300 mM NaCl, 5 mM MgCl_2_, 5 mM β-mercaptoethanol, 20 mM imidazole). The protein complex was eluted with buffers C, D and E (50 mM Tris-HCl pH 8.5, 150 mM NaCl, 5 mM MgCl_2_, 5 mM β-mercaptoethanol, 100 mM (C) / 250 mM (D) / 350 mM imidazole (E), 5 × 2 BV each). The eluted fractions were concentrated to 5 ml and loaded onto a HiLoad 26/600 Superdex 200 prep grade (GE Healthcare) equilibrated with the SEC buffer (50 mM Tris-HCl pH 8.5, 150 mM NaCl, 5 mM MgCl_2_, 2 mM β-mercaptoethanol). Confirmed by SDS-PAGE, the protein complex fractions were collected. TEV protease was added in a combination of a final concentration of 10 mM β-mercaptoethanol and incubated with gentle rocking at 4°C for approximately 12h. When the His-tag had been successfully cleaved from the protein complex, the sample was concentrated to 5 ml and loaded over the HiLoad 26/600 Superdex 200 prep grade (GE Healthcare). In order to clear the impurities which had been eluted with the complex, a reverse binding was performed by incubating for 15 minutes, rolling at 4°C with 100 μl of Ni-NTA Agarose (GE Healthcare) pre-equilibrated with the SEC buffer. The sample was separated with a gravity-flow column, and the Ni-NTA agarose was washed three times with the SEC buffer with 150 mM NaCl and finally with SEC buffer supplemented with another 150 mM NaCl (totaling 300 mM NaCl). The flow-through samples contained the clean protein complex, as verified by SDS-PAGE. The protein complex was collected, concentrated, and loaded onto the Superdex 200 Increase 10/300 GL (GE Healthcare) equilibrated with the final SEC buffer (20 mM HEPES pH 7.5, 150 mM NaCl, 5 mM MgCl_2_, 2 mM β-mercaptoethanol). The protein complex was collected and concentrated up to 5 mg/ml for further applications such as crystallization, inhibitor and activity screening. After every stage of purification, the protein content and purity were evaluated with SDS-PAGE and InstantBlue (Sigma) gel staining (Extended Data Fig. 1c).

### Protein identification from gel bands - LC-MS/MS analysis

Protein identification was performed at the Proteomics and Mass Spectrometry Core Facility, Malopolska Centre of Biotechnology, Jagiellonian University, Krakow. Samples were prepared, measured and analyzed as described in Pabis *et al*.^30^ with minor changes. Briefly, gel bands were destained by alternating washing with 25% and 50% acetonitrile (ACN) in 25 mM ammonium bicarbonate (ABC). Then, protein reduction was performed with 50 mM DTT in 25 mM ABC (45 min of incubation at 37°C) followed by alkylation with 55 mM iodoacetamide (1 h of incubation at room temperature in the dark). In the next steps, gel bands were washed with 50% ACN in 25 mM ABC, dehydrated in 100% ACN, dried and rehydrated in 20 µl of trypsin solution (10 ng/µl in 25 mM ABC). After rehydration, 40 µl of 25 mM ABC was added and samples were left for overnight incubation at 37°C. Protein digestion was stopped by adding trifluoroacetic acid (TFA) to the concentration of about 0.5%. Peptides present in the solution were collected and additionally extracted from the gel by dehydration with 100% ACN. The obtained peptide mixtures were dried and suspended in a loading buffer (2% ACN with 0.05% TFA) for LC-MS/MS analysis, carried out with a nanoHPLC (UltiMate 3000 RSLCnano System, Thermo Fisher Scientific) coupled to a Q Exactive mass spectrometer (Thermo Fisher Scientific). Peptides were loaded onto a trap column (Acclaim PepMap 100 C18, 75 μm × 20 mm, 3 μm particle, 100 Å pore size, Thermo Fisher Scientific) at a flow rate of 5 μl/min and separated on an analytical column (Acclaim PepMap RSLC C18, 75 μm × 500 mm, 2 μm particle, 100 Å pore size, Thermo Fisher Scientific) at 50°C with a 60 min gradient of ACN, from 2% to 40%, in the presence of 0.05% formic acid at a flow rate of 250 nl/min. The eluting peptides were ionized in a Digital PicoView 550 nanospray source (New Objective) and measured with Q Exactive operated in a data-dependent mode. A Top8 method was used with 35 s of dynamic exclusion. MS and MS/MS spectra were acquired with a resolution of 70’000 and 35’000, respectively. The ion accumulation times were adjusted to ensure parallel filling and detection. The acquired LC-MS/MS data were processed with the use of Proteome Discoverer platform (v.1.4; Thermo Scientific) and searched with an in-house MASCOT server (v.2.5.1; Matrix Science, London, UK) against the database of common protein contaminants (cRAP database) with manually added sequences for the proteins of interest. The following parameters were applied for the database search: enzyme: trypsin; missed cleavages: up to 1; fixed modifications: carbamidomethyl (C); variable modifications: oxidation (M); peptide mass tolerance: 10 ppm; fragment mass tolerance: 20 mmu. Additionally, the SwissProt database, restricted to *E. coli* taxonomy, was searched to assess contamination with host proteins.

### Stoichiometry determination

For protein quantitation, sample separation was carried out following a simple protocol using the Prominence HPLC system (2×LC-20AD pumps, SPD-M20A diode array detector, DGU-20 degasser, all from Shimadzu Corp., Kyoto, Japan). For gradient separation, a Kinetex 2.6 µm/100A C18 100 mm/2.1 mm ID HPLC column was used (00D-4462AN, Phenomenex, Torrance, CA, USA). Solvents used for separation: A = water + formic acid (99.9:0.1, v/v), B = acetonitrile + formic acid (99.9:0.1, v/v). All solvents were supplied by Merck local distributor (Merck KgaA, Darmstadt, Germany). Gradient was set as follows: t (time)=0 min, 25% B; t=20 min, 75% B; t=20.5 min, 90% B; t=25 min, 90% B; t=25.5 min, 25% B; t=35 min, 25% B (end). Flow rate was set to 0.3 ml/min. Diode array detector settings: wavelength range: 200-350 nm (deuterium lamp only), sampling frequency: 5 Hz. Data acquisition and data processing were controlled by LCsolution software, ver. 1.25 (Shimadzu Corp., Kyoto, Japan). To confirm protein content under every chromatographic peak taken for protein quantitation, mass spectrometry-based identification was used. The protocol for protein identification, applied with minor changes, is available elsewhere^31^. Briefly, fractions acquired during protein separation were freeze-dried using CentriVap system (Labconco, Kansas City, MO, USA) and redissolved in 70 µl 50 mM ABC (pH=7.8). Next, reduction and alkylation of cysteine residues were done using DTT and following iodoacetamide 50 mM ABC solutions (both reagents: 5 mM final concentrations). In both cases, 10 min incubation in 80°C with shaking was applied. After cooling down, trypsin (Gold-MS grade, Promega, Madison, WI, USA) was added in a final concentration of 2 pmol per sample. Samples were incubated overnight at 37°C with shaking, then freeze-dried again and redissolved in 30 µl of 4% acetonitrile/water solution acidified by 0.1% formic acid (v/v/v). NanoLC-MS/MS analyses were performed on an Ultimate 3000 system (Thermo, Waltham, MA, USA) connected on-line to AmaZon SL, equipped with nanoFlowESI ion source (Bruker-Daltonics, Bremen, Germany). Parameters of nanoLC system: column Acclaim PepMap100, C18, l=10 cm/75 µm I.D., precolumn PepMap100, C18, l=1 cm/1 mm I.D., gradient settings: solvent A = water with 0.1% formic acid (v/v), solvent B = acetonitrile with 0.1% formic acid (purity: MS-grade; Merck KGaA, Darmstadt, Germany), t (time)=0 min, 6% B; t=5 min, 6% B; t=50 min, 55% B; t=50.1 min, 80% B; t=53 min, 80% B; t=54 min, 6% B; t=58 min, 6% B (end). Sample injection volume was usually in the range 3-5 µl. Flow rate: 300 nl/min, flow rate for sample introduction on precolumn: 30 µl/min. Mass spectrometer settings were as follows: capillary voltage: 4’200V; heated capillary temperature: 150°C; MS scan range: 375-1600 m/z; MS/MS scan range: 200-2000 m/z; resolution: enhanced; scanning frequency: ca. 1 Hz; ICC (Ion Charge Control): 250’000 ions/trap cycle; fragmentation ions selection range: 450-1’600 m/z (with minor exclusions); minimal ion intensity selected for fragmentation: at least 1 × 10^6^ units. Both instruments were run under the HyStar ver. 4.1 SR1 (Bruker-Daltonics, Bremen, Germany). Data analysis was performed in Bruker’s Compass DataAnalysis 4.4 SR1. Acquired data were converted into mgf files using built-in scripts, introduced into Mascot search engine (ver. 2.6, Matrixscience, London, UK), and searched against SwissProt and in-house created database. Mascot settings: enzyme: trypsin; missing cleavages: 1; taxonomy: all; fixed modifications: carbamidomethylation; variable modifications: oxidation-Met; peptide tolerance: 1.2 Da; #13C: 1; MS/MS tolerance: 0.6 Da; peptide charge: +1,+2,+3; instrument: ion-trap.

### MicroScale Thermophoresis

His-tag proteins were labeled with Monolith His-Tag Labeling Kit RED-tris-NTA 2nd Generation according to manufacturer’s guidelines. Labeled proteins were diluted in PBS containing 0.05% Tween-20 up to 80 nM concentration and mixed with tested ligands. Samples were allowed 30 min incubation at RT prior measurements. Measurements were performed on Monolith NT.115 in duplicates using Excitation Power: 80% and MST Power: high in Monolith NT.115 Capillaries.

### NanoDSF

NanoDSF was performed in standard capillaries using Tycho equipment. Proteins and their respective complexes were measured at 1 mg/mL using default ramp temperature. The resulting melting temperatures were reported as the first derivative of the fluorescence ratio.

### Methyltransferase activity

The methyltransferase activity of wild type heterotrimer nsp10/14/16 or the heterotrimer with mutated nsp14 protein (ExoN mutant and catalytic mutant – D90A and E92A) was measured using the EPIgeneous Methyltransferase kit from Cisbio as previously described^32^. Individual kit reagents were reconstituted according to the manufacturer’s instructions. Briefly, the methyltransferase reaction was incubated for 20 minutes at room temperature in 8 µl reaction volume with 100 nM nsp10/14/16 wild type or mutated heterotrimer, 1 µM Ultrapure SAM (Cisbio), 187.5 µM RNA cap analogue (GpppA or GpppG, New England Biolabs) or 18.75 µM Cap 0 RNA oligo (TriLink) in reaction buffer consisting of 20 mM Tris-HCl pH 7.4, 150 mM NaCl, and 0.5 mM DTT. The reaction was quenched by the addition of 2 µl of 5M followed by the addition of 2 µl Detection Buffer 1 (Cisbio) to the reaction mixture. After 10 min, 4 µl of 16× SAH-d2 conjugate solution (Cisbio) was added. After 5 min, 4 µl of 1× α-SAH Tb Cryptate antibody solution was added to the reaction mixture and incubated for 1 hour at room temperature. Homogenous Time-Resolved Fluorescence (HTRF) measurements were performed on a SpectraMax iD5 plate reader (Molecular Devices) according to the manufacturer’s guideline (excitation wavelength 340 nm, emission wavelengths 665 and 620 nm, top mode, 100 flashes, optimal gain, z position calculated from the negative control (no enzyme), lag time of 60 µs and the integration time of 500 µs). The resulting data were background-subtracted and normalized as follows. The ratio of 665 to 620 nm wavelength was calculated. The data was background-corrected on the averaged signal for the buffer control. Next, the data was normalized for each series individually on the wells not containing the enzyme.

### Binding and exonuclease activity assays

Substrate for binding and exonuclease assays had a primary Sequence (5’ to 3’) of CoV-RNA1-X (X = G, or A) XXXXXXXXXXXCGCGUAGUUUUCUACGCG. The CoV-RNA1-A and G RNAs were ordered from IDT as a PAGE-purified and desalted oligos. Protein complexes (equimolar at 1.6 µM) were incubated with 100 ng of RNA in a buffer containing 20 mM HEPES pH 7.5, 100 mM NaCl, 5% v/v glycerol, 10 mM MgCl_2_, and 5 mM β-mercaptoethanol, followed by analysis via a 20% urea-denaturing PAGE or a native 1% TBE agarose gel.

### SAXS

Samples were measured in SEC-SAXS mode at BM29/ESRF, Grenoble France on 12.12.2020. (session ID MX2341). Samples (100 µL) were measured on Agilent AdvanceBio SEC 300 with 50mM Tris-HCl pH 8.5, 150mM NaCl, 5mM MgCl_2_, and 2 mM β-mercaptoethanol running phase at 0.16 ml/min flowrate. The measurements were performed at 0.99 Å wavelength. The sample to detector distance was set at 2.83 m with Pilatus2M detector for data acquisition.

### CryoEM

Out of 100 randomly chosen micrographs, 1’000 particles were manually picked without any structural knowledge about the complex to minimize bias and assigned with 2D classes that were used in the *ab initio* model built. Out of generated 3D classes, one was manually picked and used for the training of TOPAZ neural networks, which in turn picked the next interaction of particles that were used to retrain Topaz. With this approach, approximately 0.5 M particles were selected to generate 50 2D classes, out of which 19 were manually picked.

### Crosslinking and Negative-Staining Transmission Electron Microscopy

Following SEC in HEPES buffer, the purified protein complex was crosslinked with bis(sulfosuccinimidyl)suberate (BS^3^, Thermo Scientific), an amine-to-amine crosslinker. The protein sample was incubated with 0.5 mM BS^3^ (from 50 mM stock) for 30 min at RT. The crosslinking reaction was quenched with 1M Tris pH 7.5 to a final 50 mM Tris, and incubated 15 min at RT. The excess BS^3^ crosslinker was cleared by sequential dilution and concentration with Tris buffer. Concentrated samples were kept at 4°C or stored at −80°C for further measurements. Negative-stain transmission electron microscopy (NS-TEM) measurements were done in Formvar/carbon-supported 400 mesh copper grids, suspended in air with a negative lock tweezer. The purified protein complex (0.03 mg/mL) was applied on glow-discharged Formvar/carbon-supported 400 mesh copper grids and negatively stained with 1% neutralized uranyl-acetate. Grids were imaged using the JEOL JEM 2100HT electron microscope (Jeol Ltd, Tokyo, Japan) at accelerating voltage 200 kV. Images were taken by using a 4kx4k camera (TVIPS) equipped with EMMENU software ver. 4.0.9.87.

### Image processing and 3-D reconstitution

Collected micrographs were processed using cryoSPARC 3.1.1. Initially, 9’350 particles were picked from micrographs using Blob Picker. Picked particles were subjected to a template-free 2D classification, from which 1’216 particles were selected and subjected to 3D reconstitution using the ab-initio reconstitution job. The nsp10/14/16 complex map derived from SAXS data was used for a rigid-body fit in to 3D-reconstitution map using Dock in map.

## Acknowledgements

This work was supported by a subsidy from the Polish Ministry of Science and Higher Education for research on SARS-CoV-2 and a grant from the National Science Center (UMO-2017/27/B/NZ6/02488) and by EU-Horizon2020 ITN OrganoVir grant 812673 to K.P., and by the Bayerische Forschungsstiftung grant AZ-1453-20C to G.P and M.S. and a DFG Grant PO 1851/4-1 “Biochemische und strukturelle Charakterisierung des SARS-CoV-2 non-structural protein 16 (nsp16), eine cap ribose 2′O-methyltransferase.” to G.P. and M.S. The mass spectrometry was also partially financed from the subsidy no 16.16.160.557 of the Polish Ministry of Science and Education to P.S. and K.H.. J.P. is a recipient of the START fellowship from the Foundation for Polish Science (FNP). We are grateful to Local Contact at the ESRF for providing assistance in using beamline BM29. We acknowledge the MCB Structural Biology Core Facility (supported by the TEAM TECH CORE FACILITY/2017-4/6 grant from the Foundation for Polish Science) for valuable support. We acknowledge SOLARIS National Synchrotron Radiation Centre for the access to the cryo-EM Facility, where the measurements were performed.

## Extended Data

**Extended Data Fig. 1.**
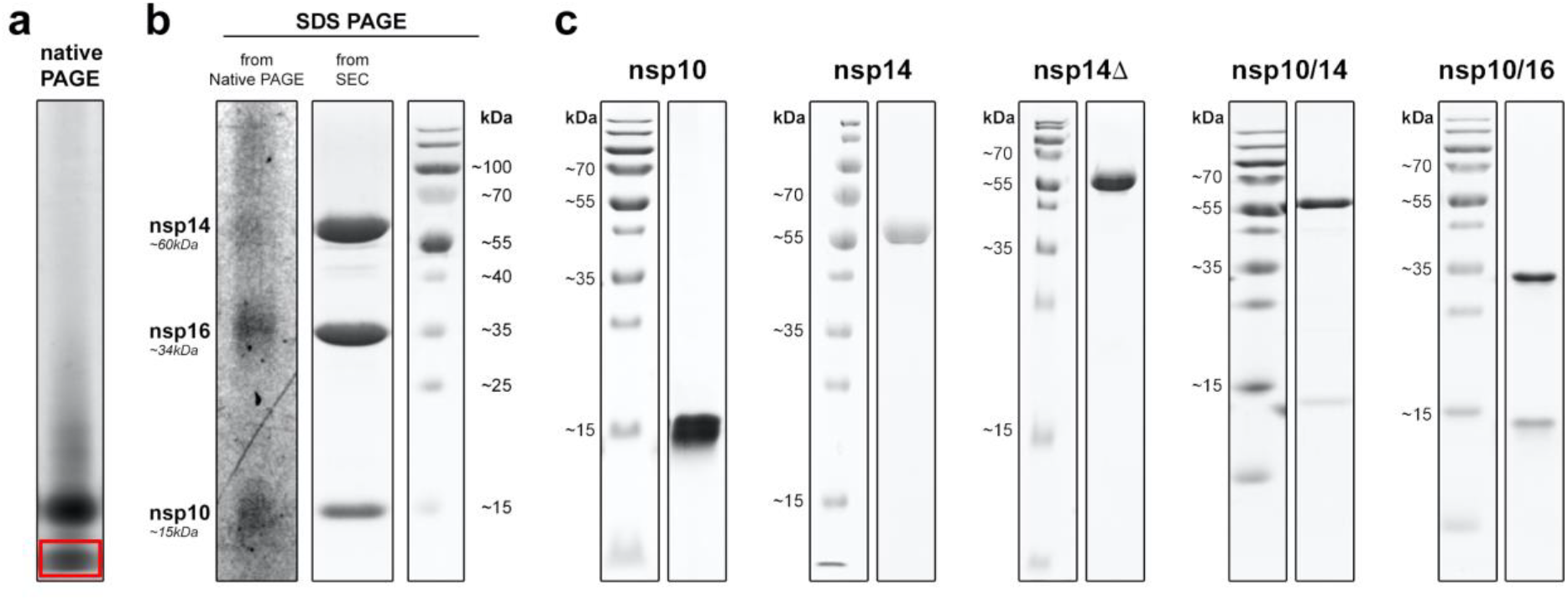
**a**, Native-PAGE of heterotrimer. **b**, SDS-PAGE of the band excited from the native-PAGE (red box showing disintegration of the heterotrimer into nsp14, nsp16, and nsp10. c, SDS-PAGE illustrating the purity of nsp10, nsp14, nsp14Δ, nsp10/14, and nsp10/16 proteins.

**Extended Table 1.**
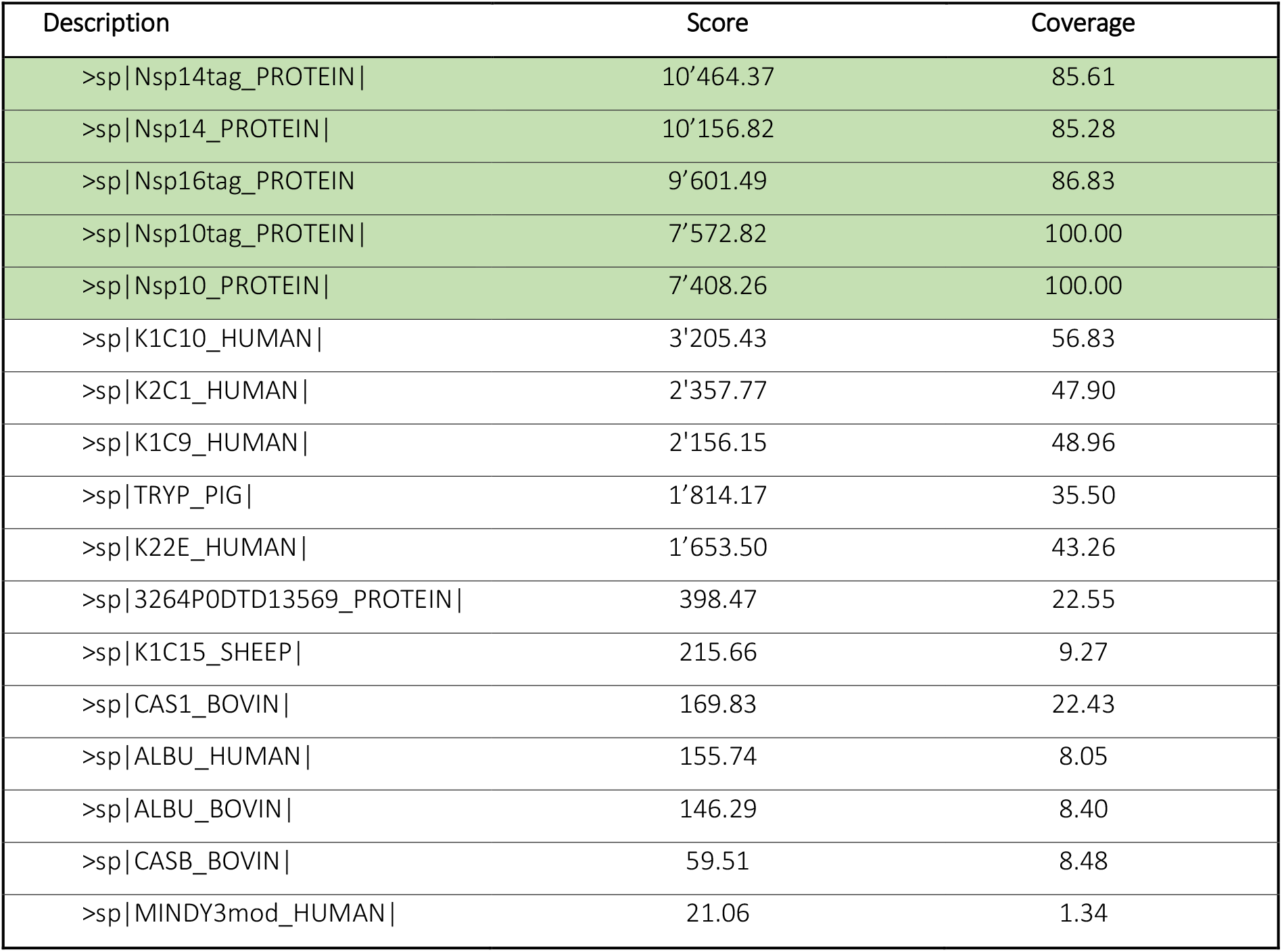
Qualitative MS analysis of the native PAGE gel band indicated in red box, in Extended Data Fig. 1a.

**Extended Data Fig. 2.**
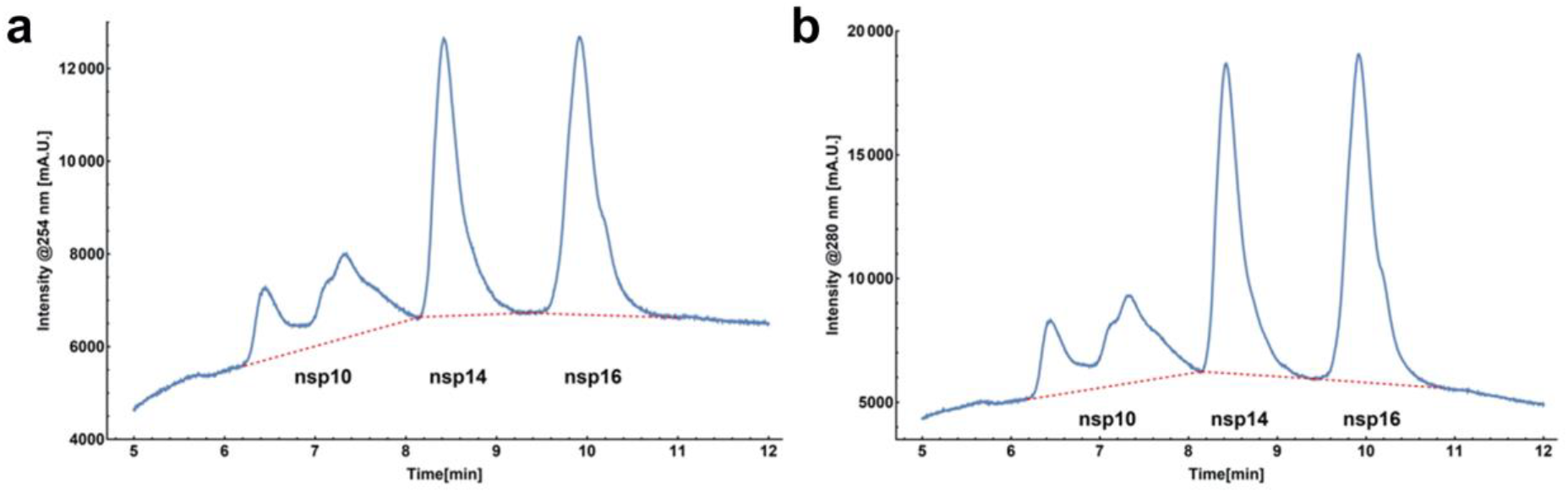
Mass spectroscopy of the heterotrimer at 254 and 280 nm.

**Extended Data Table 2.** Summary of the peak volumes from Extended Data Fig. 2 presenting 1.2:1:1 ratio of nsp10/14/16.

|  | Peak 1<br>(fr. 6.3-8.0) | Peak 2<br>(fr. 8.2-9.3) | Peak 3<br>(fr. 9.6-10.7) | Average<br>surface area | Total | Percent | Ratio |
| --- | --- | --- | --- | --- | --- | --- | --- |
| nsp10 | 64.5% (100) | 22% (19.6) | 13.5% (10.9) | 145'756 | 205'562 | 38% | 1.2 |
| nsp14 | 0% (0.0) | 90% (80.4) | 10% (8.2) | 187'293 | 167'747 | 31% | 1.0 |
| nsp16 | 0% (0.0) | 0% (0.0) | 100% (80.9) | 211'899 | 171'638 | 31% | 1.0 |

**Extended Data Fig. 3.**
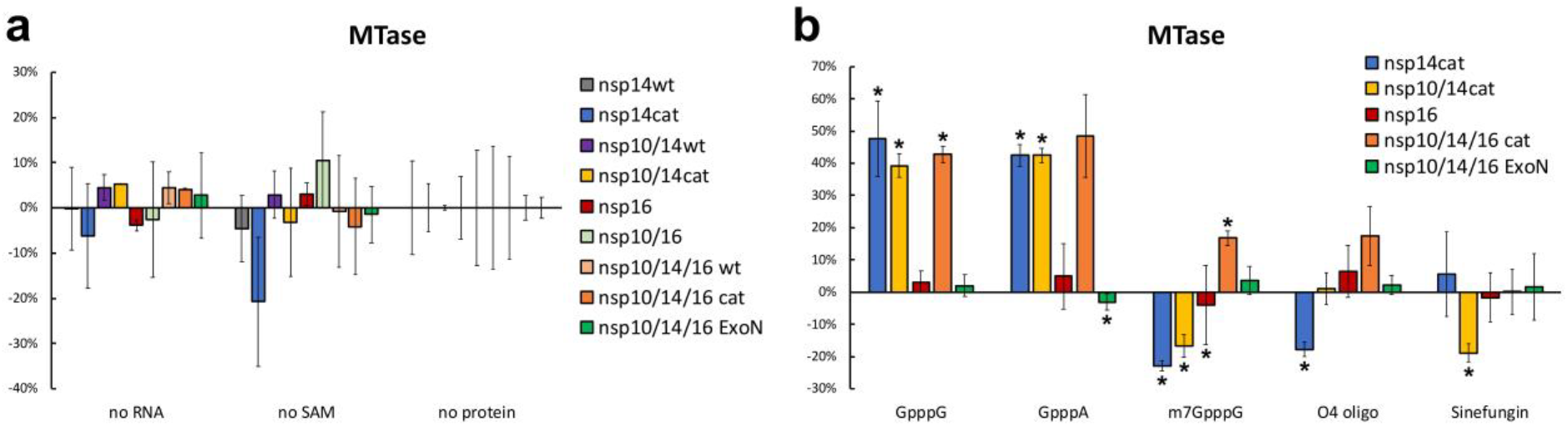
Modulation of the methyltransferase activity by the protein partners.

**Extended Data Fig. 4.**
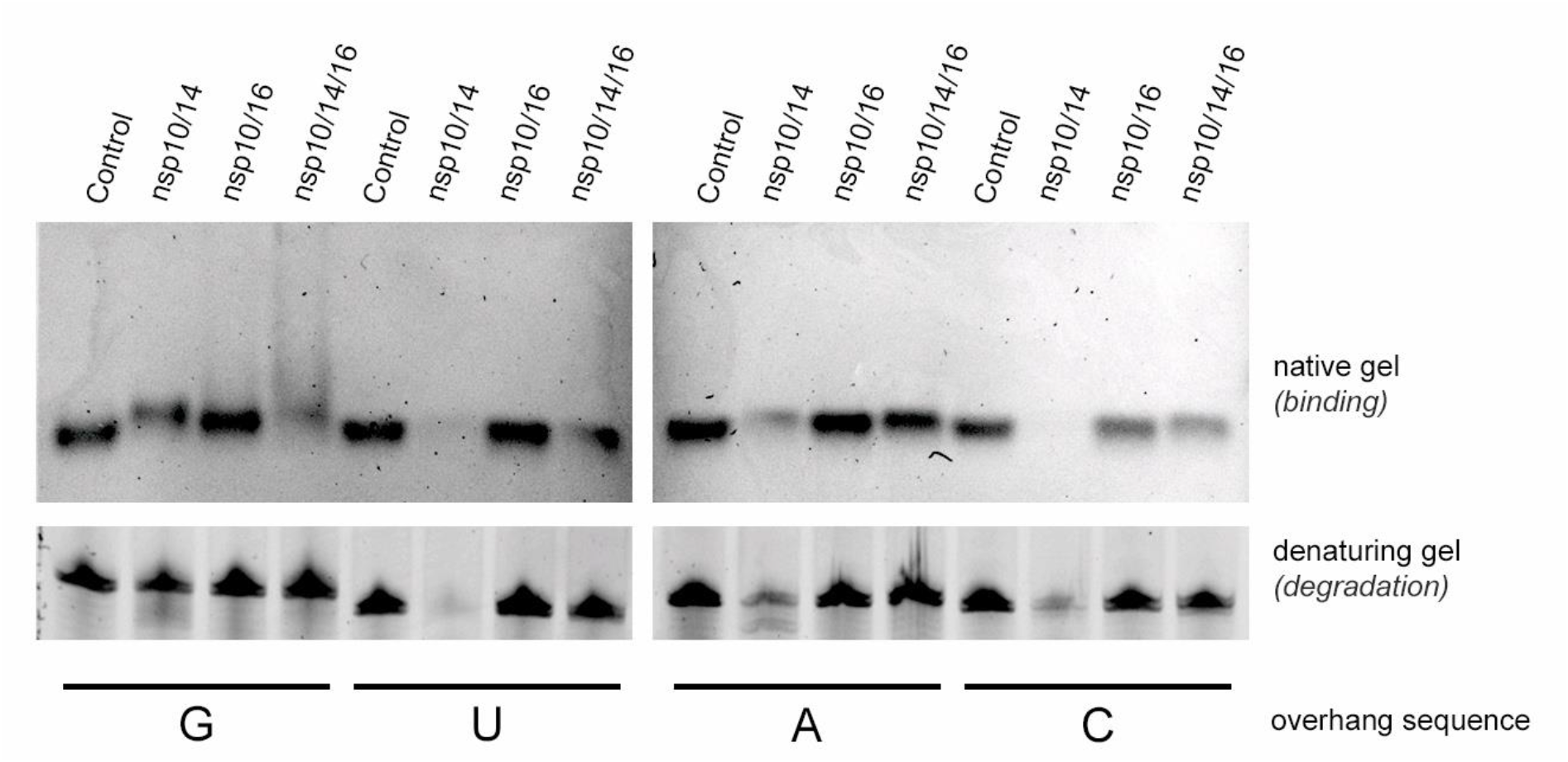
Uncropped RNA gells presented on Fig. 2.

**Extended Data Table 3.** Summary of the structural parameters derived from scattering profiles.

| a. Sample details |  |  |  |  |
| --- | --- | --- | --- | --- |
|  | nsp14 | nsp10/14 | nsp10/16 | nsp10/14/16 |
| Source organism | Severe acute respiratory syndrome coronavirus 2 |  |  |  |
| Source | E. coli BL21 |  |  |  |
| UniProt sequence ID<br>(residues in construct) | PODTD1<br>(5926-6452) | PODTD1<br>(5926-6452) | - | PODTD1<br>(5926-6452) |
|  | - | PODTC1<br>(4254-4392) | PODTC1<br>(4254-4392) | PODTC1<br>(4254-4392) |
|  | - | - | PODTD1<br>(6799-7096) | PODTD1<br>(6799-7096) |
| Molecular weight from chemical<br>composition (kDa) | 60 | 75 | 48.5 | 108.5 |
| SEC-SAXS column | AdvanceBio Bio SEC 300 column |  |  |  |
| Loading concentration (mg/mL) | 4 | 3.5 | 2.6 | 2.8 |
| Injection volume (μL) | 100 |  |  |  |
| Flow rate (mg/mL) | 0.16 |  |  |  |
| Running phase composition | 50mM Tris, 150mM NaCl, 5mM MgCl<sub>2</sub>, βME 2mM pH8.5 |  |  |  |
| b. SAXS data-collection parameters |  |  |  |  |
| Instrument | BM29, ESRF, Grenoble France |  |  |  |
| Detector | Pilatus2M |  |  |  |
| Wavelength | 0.99 Å |  |  |  |
| Sample to detector distance | 2.83 m |  |  |  |
| c. Structural parameters |  |  |  |  |
|  | nsp14 | nsp10/14 | nsp10/16 | nsp10/14/16 |
| Guinier analysis |  |  |  |  |
| I(0) (cm<sup>-1</sup>) | 8.43 ± 0.13 | 3.22 ± 0.13 | 1.73 ± 0.09 | 5.26 ± 0.11 |
| R<sub>g</sub>(Å) | 28.0 ± 0.5 | 30.1 ± 1.8 | 21.0 ± 1.5 | 40.3 ± 1.0 |
| qR<sub>g</sub>(Å<sup>-1</sup>) | 0.59 – 1.51 | 0.59 – 1.32 | 0.52 – 1.39 | 0.76 – 1.4 |
| P(r) analysis |  |  |  |  |
| R<sub>g</sub>(Å)/<br>I(0) (cm<sup>-1</sup>) | 29.4/8.61 | 32.5/3.28 | 21.1/1.64 | 41.2/5.23 |
| Guinier R<sub>g</sub>(Å)/<br>I(0) (cm<sup>-1</sup>) | 29.4/8.61 | 32.7/3.28 | 21.2/1.64 | 41.5/5.23 |
| r<sub>max</sub>(Å) | 95.8 | 122.0 | 80.0 | 140.0 |
| Total quality estimate | 0.94 | 0.78 | 0.75 | 0.77 |
| Molecular weight estimate/predicted<br>(kDa) | 33.1/60 | 28.9/75 | 20.6/48.5 | 83.2/108.5 |
| Oligomerization state | monomeric |  |  |  |
| d. Shape model-fitting results |  |  |  |  |
| DAMMIF (10 runs) | nsp14 | nsp10/14 | nsp10/16 | nsp10/14/16 |
| q<sub>max</sub> range for fitting(Å<sup>-1</sup>) | 0.26 | 0.26 | 0.21 | 0.17 |
| Symmetry, anisotropy assumptions | P1, none |  |  |  |
| NSD (standard deviation) | 1.42 (0.07) | 1.10 (0.07) | 1.29 (0.14) | 0.80 (0.08) |
| Chi-squared | 1.16 | 1.09 | 1.07 | 1.08 |
| Resolution (from SASRES) (Å) | 43 ± 3 | 39.3 ± 3 | 33.3 ± 3 | 38.3 ± 3 |
| SASDBD IDs | SASDKT6 | SASDKU6 | SASDKV6 | SASDKW6 |
| e. Oligomer volume fractions |  |  |  |  |
| volume fractions Chi^2 1.06 |  |  |  |  |
| nsp10/16 (6YZ1) | nsp16 | nsp14 | nsp10 | nsp10/14 (5CU8U) |
| 53% | 0% | 47% | 0% | 0% |

**Extended Data Fig. 5.**
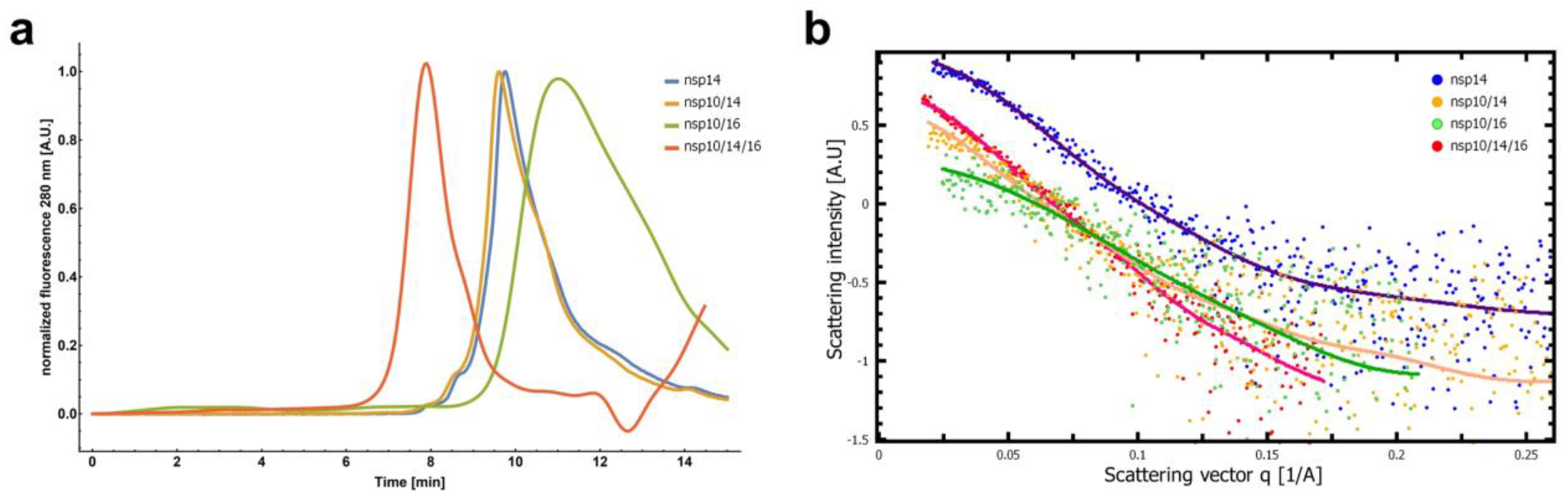
SEC-SAXS. **a**, The SEC profiles for nsp10/14/16 (red), nsp10/14 (yellow), nsp14 (blue) and nsp10/16 (green) directly before flow cell for SAXS measurement. The 280 nm fluorescence intensity was normalized for clarity. **b**, SAXS scattering profiles resulting from merging the signal form SEC-SAXS experiments for nsp10/14/16 (red), nsp10/14 (yellow), nsp14 (blue) and nsp10/16 (green). The solid lines represents the fit of the experimental data to the real space models. In both cases the position closer to the left (lower elution volumes or the curvature change at smaller scattering vector values q) indicate larger objects that were analyzed. Therefore, nsp10/14/16 represents the largest from the analyzed protein complexes, while nsp10/16 the smallest, which is in the agreement with their theoretical masses calculated from the amino acid sequences.

**Extended Data Fig. 6.**
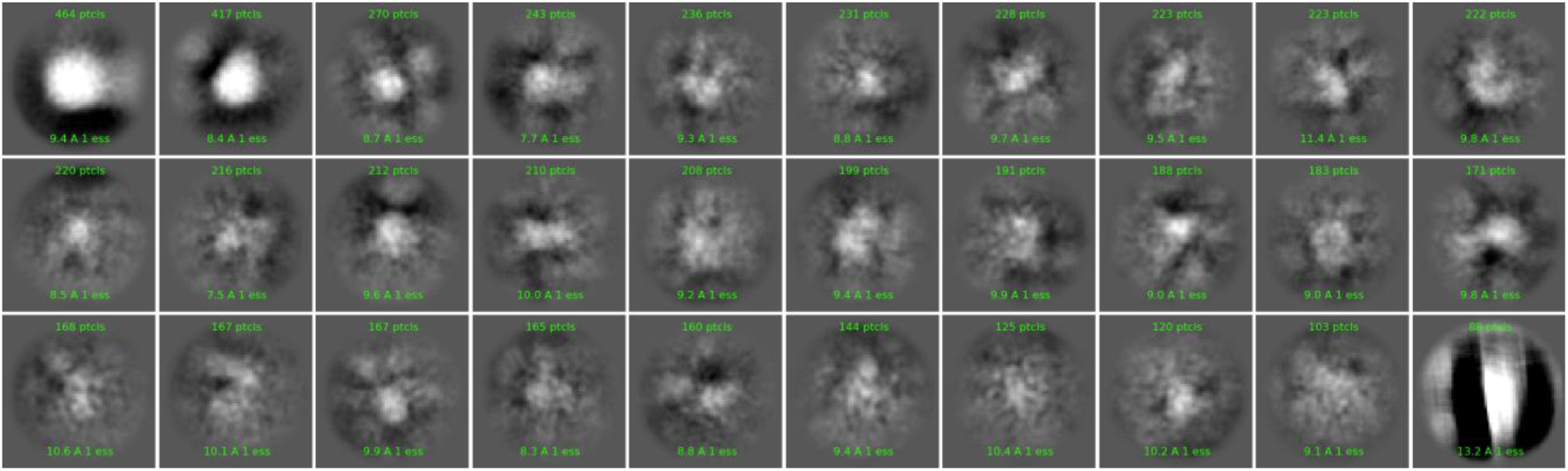
2D classes picked in cryoSparc for full length heterotrimer.

### CryoEM

We attempted to solve a high-resolution heterotrimer structure using cryoEM techniques. Highly purified heterotrimer solution was vitrified on grids using Vitrobot under various conditions. Resulting grids were measured using Titan Krios G3i at Solaris, Poland. Data analysis was performed using cryoSPARC software. We managed to train the TOPAZ neural networks to pick approximately 0.5 M particles used to generate 50 2D classes, out of which 20 were manually picked. Generated classes, though noisy, exhibit croissant-like shape with four distinct domains that correspond to nsp16, nsp10 and two domains of nsp14 (highlighted with arrows on the class that shows side-on view). Unfortunately, we were not able to reconstruct a high-resolution 3D structure from collected data. Our best attempt shown below in cyan has ca. 9 Å resolution. The fitted heterotrimer hybrid model based on SAXS data (gold) shows overall good fit with the extra volume that may arise from the flexible nature of the heterotrimer.

**Extended Data Fig. 7.**
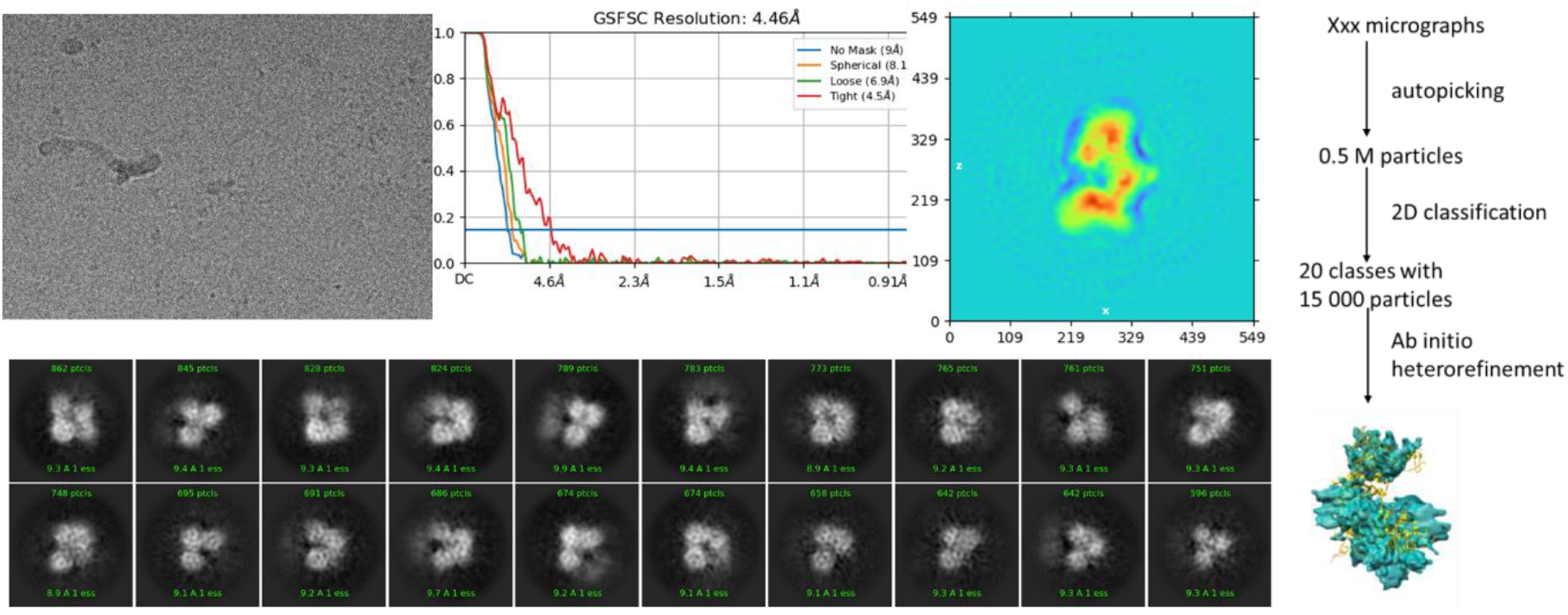
An overview of cryoEM data analysis in cryoSPARC. The overlay of generated 3D model from cryoEM (cyan) and SAXS model (gold).

